# Development of an absolute assignment predictor for triple-negative breast cancer subtyping using machine learning approaches

**DOI:** 10.1101/2020.06.02.129544

**Authors:** Fadoua Ben Azzouz, Bertrand Michel, Hamza Lasla, Wilfried Gouraud, Anne-Flore François, Fabien Girka, Théo Lecointre, Catherine Guérin-Charbonnel, Philippe P. Juin, Mario Campone, Pascal Jézéquel

## Abstract

Triple-negative breast cancer (TNBC) heterogeneity represents one of the main impediment to precision medicine for this disease. Recent concordant transcriptomics studies have shown that TNBC could be splitted into at least three subtypes with potential therapeutic implications. Although, a few studies have been done to predict **TNBC subtype** by means of **transcriptomics data**, subtyping was partially sensitive and limited by batch effect and dependence to a given dataset, which may penalize the switch to routine diagnostic testing. Therefore, we sought to build an absolute predictor (i.e. intra-patient diagnosis) based on **machine learning** algorithm with a limited number of probes. To this end, we started by introducing probe binary comparison for each patient (indicators). We based predictive analysis on this transformed data. Probe selection was first performed by combining both filter and wrapper methods for **variable selection** using cross validation. We thus tested three **prediction models** (random forest, gradient boosting [GB] and extreme gradient boosting) using this optimal subset of indicators as inputs. Nested cross-validation allowed us to consistently choose the best model. Results showed that the 50 selected indicators highlighted biological characteristics associated with each TNBC subtype. The GB based on this subset of indicators has better performances as compared to the other models.

## Introduction

Triple-negative breast cancer (TNBC), which lacks the expression of estrogen receptor, progesterone receptor, and HER2, accounts for 12 to 17% of primary breast cancers, and is one of the most aggressive and deadly breast cancer subtypes [1]. Furthermore, heterogeneity and lack of targeted therapies represent the two main issues for precision treatment of TNBC patients. Molecular subtyping and identification of therapeutic pathways are therefore required to optimize medical care of these patients. Recently, we identified three TNBC subtypes with potential therapeutic implications: molecular apocrine (C1) and two basal-like enriched with opposite immune responses (C2: immunosuppressive response; C3: adaptive immune response) [2–4]. These results were based on transcriptomic data (Affymetrix® Human Genome U133 Plus 2.0 Arrays) and validated by means of external data and protein expression using immunohistochemistry and proteomic data. Therefore, transition to precision medicine and selection of targeted treatments for TNBC depends on the establishment of a TNBC subtype predictor.

The aim of the present work was to build such a predictor able to classify new patients by means of an algorithm that uses a set of Affymetrix® probeset intensities. In order to optimize the robustness of the algorithm and the likely manufacture of a diagnostic test, the following methodological guidelines have been imposed: Absolute assignment of TNBC subtypes, independent of external data (“intra-patient diagnosis” or “universal predictor”), and limited number gene probes, respectively. The first requirement was made to avoid the need for external data and/or normalization, and to circumvent molecular subtyping shortcomings [5–8]. The second requirement took into account the optimization of the diagnostic test from an industrial point of view, implying that it is preferable to select a low number of gene probes.

The identification of TNBC subtypes can be seen as a classification task, which can be addressed with machine learning methods. In the healthcare field, these techniques are popular and even essential to analyze the large volumes of data produced for a patient. In fact, the healthcare data generated are sometimes too complex and voluminous to be processed by traditional methods. Analytical methods then provide tools to turn this data into useful information for decision-making and useful knowledge for biomedical research [9,10].

Our main objective was to build a predictive model based on the observation of 54.675 probes for providing the probabilities of belonging to each TNBC subtypes for each patient. Note that this objective is different and more tricky than only predicting the TNBC subtype. Probability estimation for classification has a long-standing tradition in biostatistics and machine learning, see for instance Kruppa et al. [11,12]. For our issue, this strategy was justified not only by the fact that our targets represent a membership value for each patient belonging to each of the TNBC subtype (one score by subtype), but also, by the existence of biological gradients between the two basal-like TNBC subtypes (C2 and C3). Two biological characteristics stood out: immune response and *PIK3CA* mutation status. Concerning the first one, two opposite immune response gradients were observed. Archetypal C2 tumors displayed the highest pro-tumorigenic response (high M2/M1 ratio), decreasing steadily from C2 to C3. To the contrary, archetypal C3 tumors displayed the highest anti-tumorigenic responses (T-cell, B-cell, T-cell cytolytic activity, STAT1, MHC-2, Type I interferon, MHC-1), decreasing steadily from C3 to C2. The *PIK3CA* mutation status gradient showed a profile comparable to that observed for anti-tumorigenic ones. C2 and C3 tumors have a common basal-like background but mainly differ by the direction and the level of the immune response, and the level of PIK3CA mutation status. Probabilities were very close in the border between C2 and C3 and can lead to ambiguity in group memberships. It is thus of first importance to estimate these probabilities.

Our prediction method takes as an input a given fuzzy clustering and more specifically the probabilities of belonging to each cluster. In the paper, we illustrate the method on the results of Jézéquel et al. which provide a fuzzy clustering of the TNBC subtypes and the corresponding TNBC subtype probabilities over 693 patients [4]. The problem then boils down to a multi target regression methodology based on 54.675 probes and can be solved by various methods in machine learning. Note that the method could be applied to the results of other clustering algorithms of TNBC and in particular with more clusters.

There are two distinct reasons to apply variable selection for reducing the number of variables (probes): first because from the industrial point of view we want a low number of gene probes and corresponding measures (the parsimony principle), second because we want to control the overfitting of models fitted on the data. After variable selection, three machine learning methods were applied: 1) random forest (RF); 2) gradient boosting (GB); 3) XGBoost (XGB) (extreme gradient boosting) [13–15]. The three models have many parameters which can be optimized in order to improve model’s performance. The best model was then selected: Cross validation (CV) for both model selection and tuning parameters, however two different CV are required. Thus, nested CV (NCV) was performed to get an unbiased estimate of performance of predictive model [16–18].

## Materials and methods

### TNBC cohorts

We looked for TNBC cohorts with available genomic data. To avoid cross-platform normalization issues, we exclusively looked for Affymetrix® genomic datasets using the same platform, here the HG U133Plus2 microarray, in repositories such as Gene Expression Omnibus (GEO) and ArrayExpress, selecting those with a medium to large sample size. All datasets were MAS5-normalized using the Affymetrix Expression Console software with default analysis parameters and then log2-transformed.

### Experimental design

We aimed at identifying TNBC subtypes and to establish an universal predictor able to predict the subtype of a new patient. To do this, we carried out this study in two steps. In the first step, clustering analysis was conducted based on several transcriptomic datasets. We have been careful to remove batch effect before unsupervised analysis. In the second step, we selected variables to include to predictive models (Figure 1). We built several predictive models and then chose the best one able to predict the TNBC subtype of new patients.

**Figure 1.**
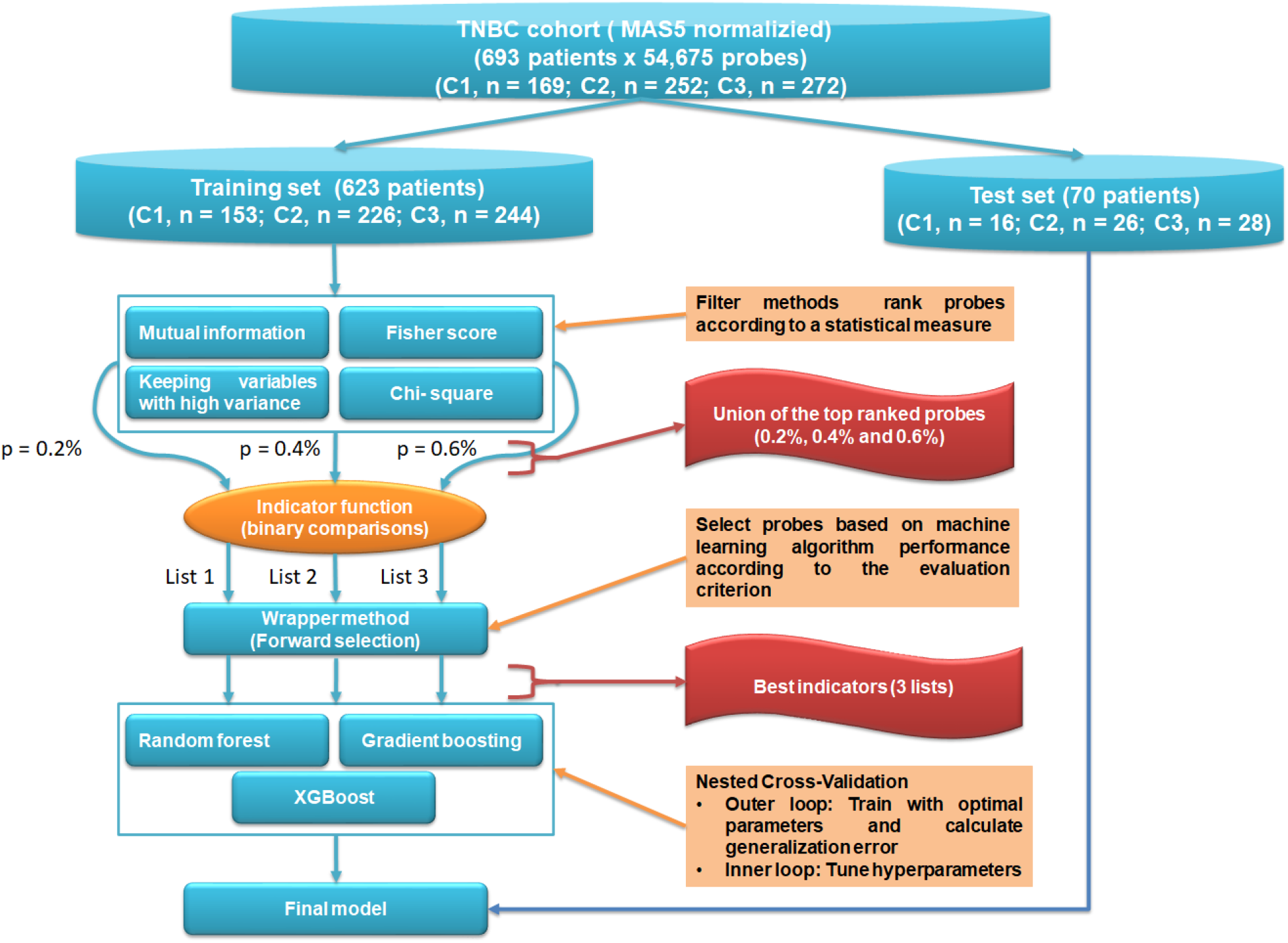
Flowchart of subtype prediction and variable selection. The study can be divided into two mains steps: variable selection and model selection and evaluation. For the optimal variable selection, several univariate filter methods were used to select preliminary probes subset, which was then used as input for the wrapper method (Forward selection) after binarization using indicator function. Prediction subtype using several models with the optimal selected indicators was applied. Nested cross-validation was used to train and evaluate models. Based on the optimal subsets of indicators, the best predictive model was applied to the test set after binarization.

### Cluster analysis to identify subtypes of TNBC

#### Data integration and batch effect removal

All cohorts were merged into a single one to produce an information-rich dataset and hence to increase the quality and performance of statistical analysis. However, the main problem of microarray data integration is batch effect due to different studies. Many methods exist for removing batch effects from data [19,20]. We tested two of the most used methods: “removeBatchEffect” (RBM) in the “limma” package and “Combat” [21,22]. Guided principal component analysis (gPCA) was used to evaluate them and then to choose the best one for our dataset [23].

#### Class partition

Fuzzy C-means (FCM) clustering was used to cluster data into homogeneous subgroups with similar biological characteristics [2,4,24]. We performed FCM clustering with centered Pearson distance based on the 5% most variable probe sets (n = 2,734 probes) after batch effect removal. FCM provides for each patient the corresponding probabilities of the TNBC subtypes.

### Subtypes prediction and variable selection

For improving the robustness of our method, we built prediction rules on comparison variables between all pairs of probes, rather than on the raw variables (MAS5 normalized data) by introducing comparison variables. These binary comparisons have been named “indicators”. The indicator “A > B” were equal to “1” if the expression value of probe A was greater than the expression value of probe B, and “0” otherwise. In consequence, analyses were less sensitive to standardization methods or batch effect removal methods [25–28]. This approach met our first requirement. However, it increased the number of new variables (indicators) significantly and consequently increased both training time and risk of overfitting. So, in order to optimize machine learning models and to reduce the numbers of variables, we performed variable selection methods. This last point met the requirement of industrial application (that is, a “reasonable” number of probes).

#### Variable selection

The selection of variables is an important step of model design, especially when the number of variables is large. The aim of this step was to determine an optimal subset of descriptors to decrease model complexity and increase prediction score.

Here, we combined two approaches: first, univariate filter methods to reduce variable space by selecting a subset of probes regardless of the model, and then wrapper based on this subset by training a learning model that determined the usefulness of this subset and selected an optimal sub-subset according to the evaluation criterion, which depends on the type of problem. Here it’s a regression task.

Four filter approaches based on univariate statistical tests were applied to the original dataset MAS5 normalized: mutual information, Fisher score test, Chi-2 test and keeping probes with high variance (Figure 1). We used “SelectPercentile” function from “sklearn.variable_selection” Python module to select probes according to a percentile of the highest scores. Three arbitrary percentile thresholds were used to select the number of probes: 0.2% (List 1), 0.4% (List 2) and 0.6% (List 3). And then we took the union of the probes obtained by the four filter methods, and this for each percentile. In the second step, we further applied forward selection (FS) to subset of probes, which have been selected previously in the first step. FS process, explained by pseudo-code, is detailed in Figure 2. Briefly, it works in two steps. In the first step, the algorithm seeks for the best two probes to minimize the cross validation mean squared error (RMSE CV) at each step using “RandomForestRegressor” function from “sklearn.ensemble” Python module. In the second step, FS process tests the rest of the available probes. For each probe, the binary comparison indicators were built from the probes already chosen and a new probe. The CV error obtained was then calculated using old and new indicators. This step was repeated until the RMSE no longer decreased or until maximum number of probes was obtained, which was fixed beforehand (input parameter of our FS function) (Figure 2).

**Figure 2.**
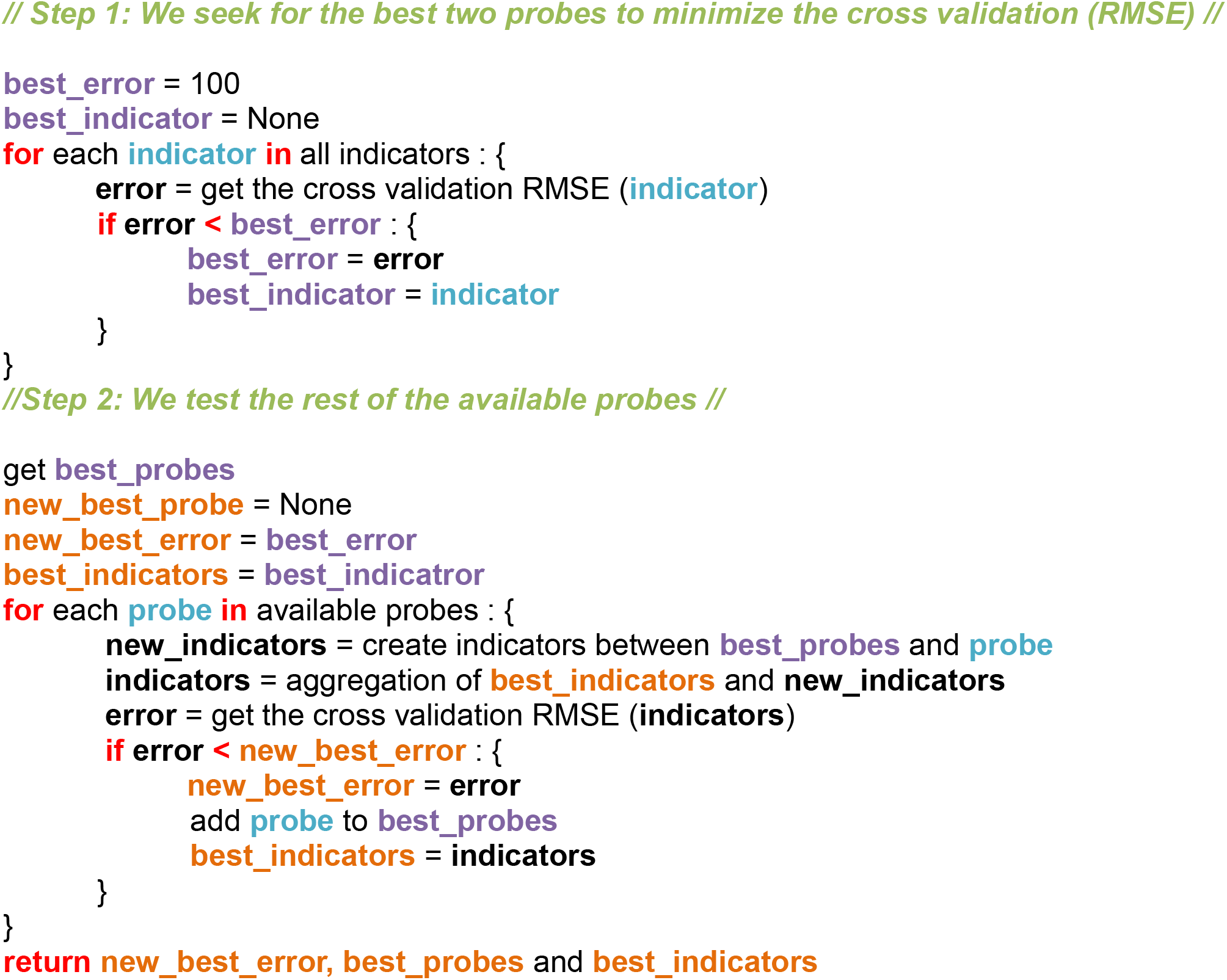
Forward selection explained by pseudo-code for identifying probes and indicators.

FS algorithm was also used to select the best indicators once the probes have been fixed.

#### Probability estimation and class prediction

Many supervised machine learning models can be used for multi target regression, that is to predict a d-dimensional vector of real values given a set of predictors (variables). We identified three of the most popular used methods to be tested in our study: 1) RF; 2) GB; 3) XGB [13–15]. We note that only RF satisfies the constraint, which guarantees the sum of membership values for each patient is equal to 1. It was therefore necessary to standardize the predictions output for GB and XGB. Two methods were tested. In the first one, we divided each of the values by the sum of the three probabilities obtained. We will use the terms GB1 and XGB1 to denote GB and XGB after this standardization, respectively. In the second one, we kept the greatest probability value and divide the two others so that the sum of the new probabilities makes one (GB2 and XGB2).

Prediction for the TNBC was also derived for each patient by selecting the TNBC class of highest probability.

#### Model selection and evaluation

To get an unbiased estimate performance of our predictive model, we split the dataset into a 90% training set and 10% test set. Machine learning models were fitted on the training data with nested cross validation (NCV). The nested CV has the inner loop CV nested in an outer loop CV. The CV inner loop is used to perform hyperparameter tuning (here executed by “GridSearchCV” from Python module “sklearn.model_selection”), while the outer CV is used to estimate the unbiased generalization performance of models. The complete workflow using stratified NCV is described in Figure 1. The cycle of nested CV was independently run ten times in order to obtain a reliable result. The metric used to evaluate prediction errors on regression models is R squared (R2) which represents the coefficient of adequacy of the values compared to the values of origin. Here, the R2 score for each individual target was averaged. We used the default “uniform averaged”, which specifies an uniformly weighted mean over outputs.

In addition, since multi-outputs represent the probability of belonging to each TNBC subtypes, we assigned each patient to the group to which he had the highest probability of belonging. Then we calculated the rate of well classified, the sensitivity and specificity of each subtype.

### Statistical analysis

Clustering and figures’ generations were performed using R (R Core Team 2018) version 3.6.1 for Windows® [29] and the packages “amap” 0.8.16, “cluster” 2.0.7 and “ggplot2” 3.3.0 packages. The subtypes prediction and variable selection statistical analysis was performed in python version 3.7.3 in Jupyter notebooks version 6.0.0 with modules like “SKLearn” 0.21.2 [30], “XGBoost” 0.90 and other functions that we wrote ourselves.

## Results

### TNBC cohorts

Seven cohorts were selected, including a total of 693 TNBC patients (Table S1).

### Unsupervised analysis

Two methods of removing batch effects were tested: RBM and Combat. To evaluate batch effect removal efficacy of each method, gPCA was used. Test statistic δ for Combat method was very close to zero compared to RBM (δ = 0.0861; P = 0.953 and δ = 0.2743; P = 0.1 for RBM, respectively). This result allowed us to choose Combat method to remove batch effects, exclude the hypothesis of technical biases and permits us to merge all cohorts. Principal component analysis (PCA) with the projection of the different cohorts onto the first principal plane showed a homogeneous distribution of patients and that there is no apparent variability related to different studies (Figure S1).

Biological significance of the clusters found in our previous studies lead us to choose the number of three clusters by means of FCM: C1 (n = 169; 24.4%); C2 (n = 252; 36.4%); C3 (n = 272; 39.2%) [2,4] (Figure S2).

### Variable selection

Table 1 summarizes and shows the number of probes and indicators selected after each step of our pipeline.

**Table 1.** Summary of the number of probes and indicators in function of the three probe threshold filter lists. List 1: 0.2%; List 2: 0.4%; List 3: 0.6%.

|  | **List 1** | **List 2** | **List3** |
| --- | --- | --- | --- |
| Threshold percentile | 0.2% | 0.4% | 0.6% |
| Probes generated per univariate filter method | 109 | 219 | 328 |
| Union of the four univariate filter methods (probes) | 361 | 675 | 975 |
| FS based on probes | 32 | 36 | 34 |
| Indicators generated | 496 | 630 | 561 |
| FS based on indicators | 50 | 50 | 50 |

Three probe lists were obtained containing 361, 675 and 975 probes for the three arbitrary threshold filters, 0.2%, 0.4% and 0.6%, respectively. Each of them represents the union of the top ranked probes generated by the four univariate filter methods (Figure 1). In the following, for convenience purposes only, we will use the terms “List 1”, “List 2” and “List 3” for each calculation step according to 0.2%, 0.4% and 0.6% threshold filters, respectively.

To visualize the results of the FS applied to the three lists of probes, we plotted the RMSE CV as a function of the number of probes in the model (Figure 3A, C and E). The three curves decreased first and then remained stable between 30 and 50 probes. We selected 32, 36 and 34 probes for List 1, List 2 and List 3, respectively (Table S2). Indeed, for List 2 and 3, the number of selected probes corresponded to the minimum value of RMSECV. However, for the List 1, the smallest RMSECV (0.1108) value was obtained when the number of probes was equal to 41, which generated 820 indicators. This number was relatively high according to our criteria. However, with 32 probes we also got a value close to the smallest RMSECV (0.1134), and we reduced the number of indicators by almost half (496 indicators). Selected probes were linked to cluster biological characteristics: molecular apocrine for C1, basal-like for C2 and C3, and immune response for C3 (Table 2).

**Figure 3.**
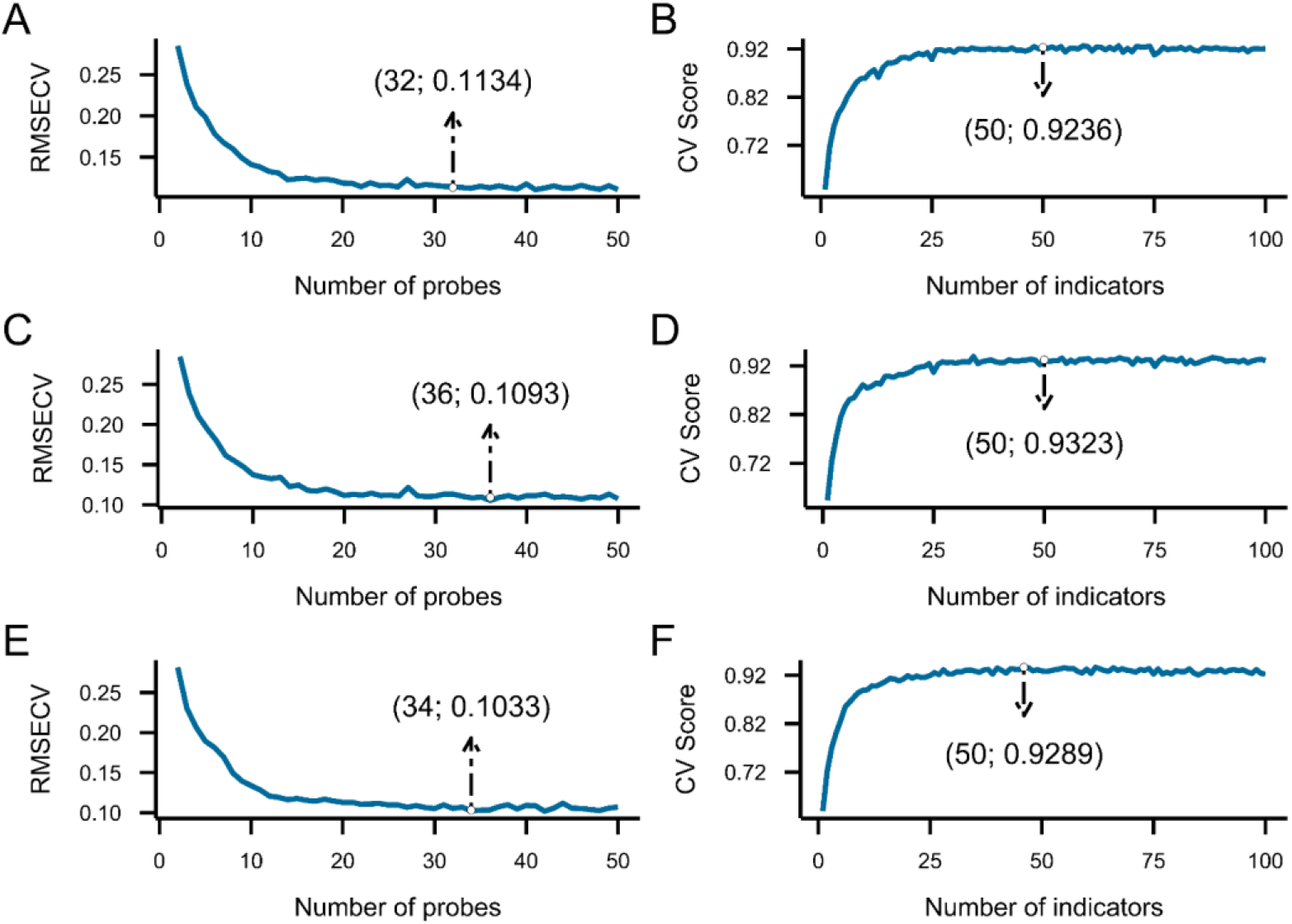
Forward selection on the probes preselected by univariate filter methods. The variation of RMSECV with the number of probes selected by forward selection on the probes using List 1 (A), List 2 C) and List 3 (E). Score depending on the number of indicators selected by forward selection based on the probes selected previously using List 1 (B), List 2 (D) and List 3 (F).

**Table 2.** TNBC subtype specificity (C1, C2, C3) of selected probes/genes in function of the three probe threshold filter lists. List 1: 0.2%; List 2: 0.4%; List 3: 0.6%.

| **List 1 (n = 32)** | **List 2 (n = 36)** | **List 3 (n = 34)** |
| --- | --- | --- |
| C1 (n = 9):<br>*AGR2; ALCAM; AR; FOXA1; ITGB5; NQO1; PRICKLE2; SMIM14; TTC8* | C1 (n = 9):<br>*ADH1B; APBB2; ALCAM; AR; DUSP4; FOXA1; KIAA1324; SLC7A8; UGDH* | C1 (n = 7):<br>*APBB2; AR; ERBB4; FAM214A; FOXA1; KIAA1324; MEGF9* |
| C2 and C2/C3 (n = 11):<br>*BCL11A (x2); CDCA2; CDCA7; COL2A1; FOXC1; KNCK5; 220425_x_at; TAF4B; TFCP2L1; TTK* | C2 and C2/C3 (n = 13):<br>*ACTG2; BCL11A (x2); CDCA2; CDCA7; FOXC1; IQCG; KRT23; MIA; PIMREG; SHC4; TTK; TFCP2L1* | C2 and C2/C3 (n = 15):<br>*ABCC11; ACTB; BCL11A; CCNE1; CENPF; FOXC1; MAP2; PIMREG; PRKX; SHC4; SOX6; SOX10; SUV39H2; TFCP2L1; YEATS2* |
| C3 (n = 12):<br>*CD3D; CYTIP; GZMA; 214916_x_at; IGKC; 216401_x_at; IKZF1; JCHAIN; NKG7; P2RY10; POU2AF1; SLA2* | C3 (n = 14):<br>*APOBEC3G; CD2; CD38; CD3D; CD52; CYSLTR1; 216401_x_at; IGLV2-14; JCHAIN; NKG7; POU2AF1; PVRIG; RHOH; SAMSN1* | C3 (n = 11):<br>*217157_x_at; 217281_x_at; 217480_x_at; AIM2; CD2; GIMAP4; IFIH1; IGLL3P; IL10RA; POU2AF1; SLAMF7* |
|  |  | Unknown (n = 1):<br>*200099_s_at* |

We proceeded in the same way for FS on the 496, 630 and 561 indicators formed by the previously selected probes for List 1, List 2 and List 3, respectively (Table 1). The relation between the number of indicators used for the construction of the model and the corresponding performances measured by R2 score obtained from CV are displayed in Figure 3B, D and F. A considerable increase in score CV was observed, but ultimately it was generally flat and stabilized with just over 30 indicators, indicating that an appropriate number of indicators was reached. We restricted the number of indicators to 50 for each of the three lists (Table S3). Overall, all indicators contributed relevant information to discriminate the three TNBC subtypes (Figure S3, S4 and S5). Each barplot highlighted a significant difference in the distribution of the TNBC subtypes between the indicator values. Taking for example the indicator “FOXA1 > TTK”, it took the value “1” at high frequency in C1 while relatively low in C2 or C3. This initial impression was confirmed by Fisher’s exact statistical test (P < 0.001). All selected indicators can be classified into four categories based on cluster separation information: C1 versus C2/C3; C2 versus C1/3; C3 versus C1/C2; C1 versus C2 versus C3 (Table 3). Distributions of the numbers of indicators between the four categories was similar, as performed by the Fisher’s exact test (P = 0.6642).

**Table 3.**
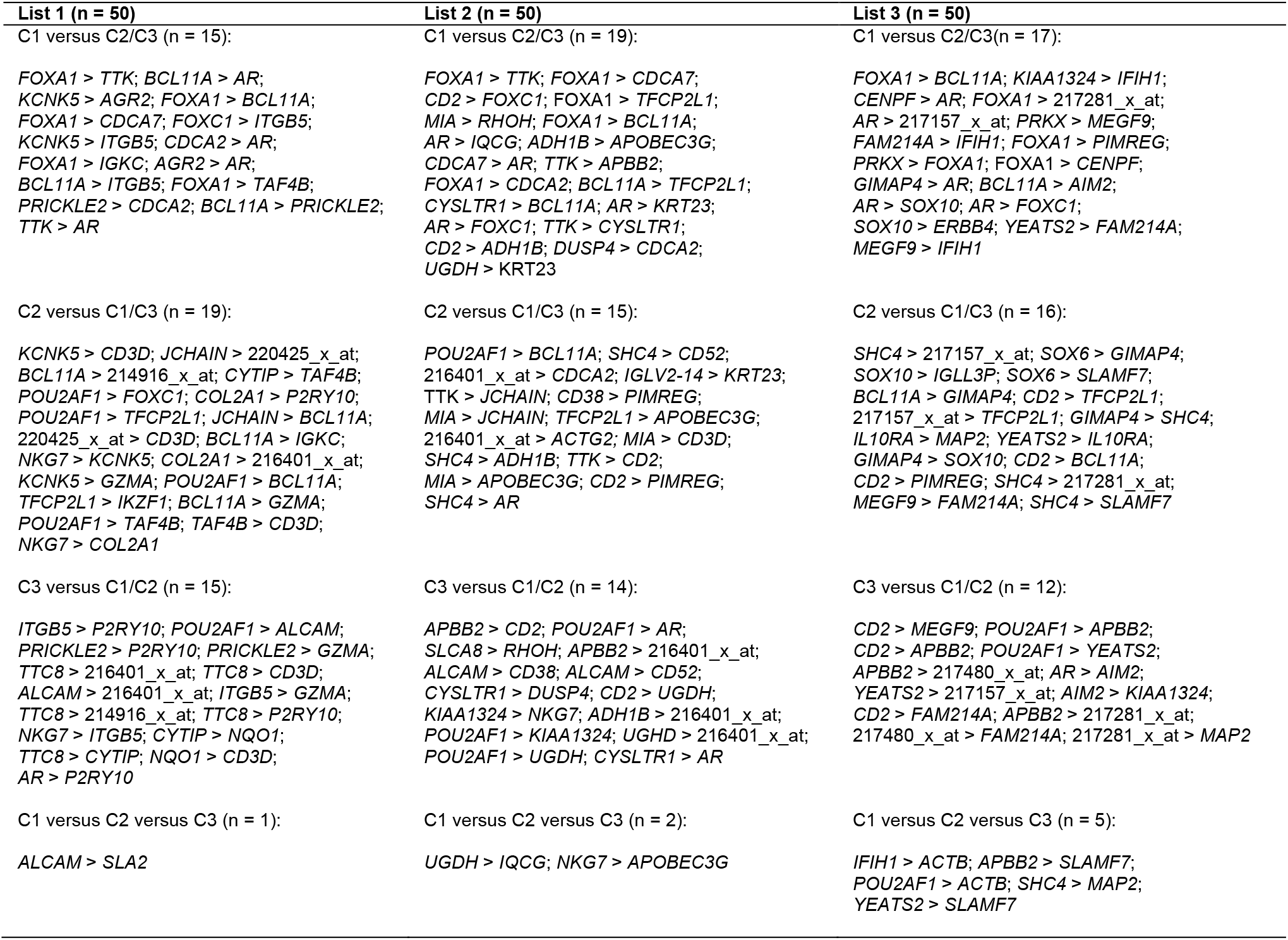
TNBC subtype specificity (C1, C2, C3) of selected indicators in function of the three probe threshold filter lists. List 1: 0.2%; List 2: 0.4%; List 3: 0.6%.

### Probability estimation and class prediction

The evaluation metrics are reported in the two boxplots Figure 4A and B, which show the average R2 score (i.e. inference of the probabilities) and average accuracy (i.e. classification into TNBC subtypes) over NCV over 10 runs, respectively. The predictive score and accuracy across the five models were consistently high. This is valid for the three lists. However, List 2 showed the best performances, both in terms of R2 scores and accuracy regardless the models. Results are summarized in Table S4 and S5.

**Figure 4.**
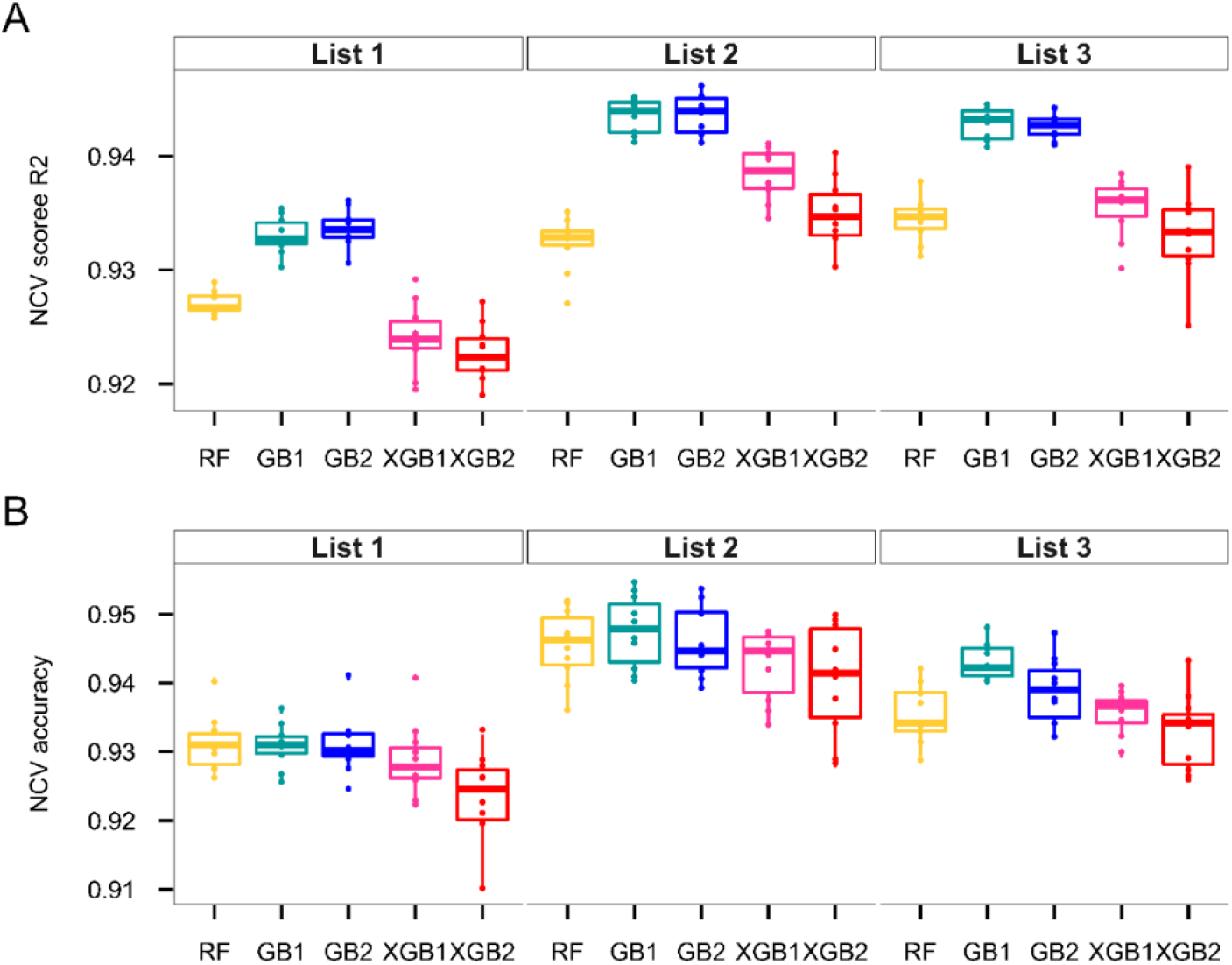
Performance comparisons of different models using different list of indicators and 10 runs of NCV. A. Boxplots showing the distribution of average R2 scores; B. Boxplots showing the distribution of average accuracy. Random Forest (RF) (yellow), Gradient Boosting 1 (GB1) (green), Gradient Boosting 2 (GB2) (blue), XGBoost 1 (XGB1) (pink) and XGBoost 2 (XGB2) (red).

For this list of indicators (List 2), the two predictive models based on gradient boosting contributed the best. They showed similar performances. So, we chose to aggregate the results of these two models instead of selecting one. We take the average of the probabilities obtained by each GB model.

The aggregation of the two models gave a slightly higher R2 score and built a compromise between the two models regarding the accuracy (Figure 5). This model was worthy of further study, it was the best and was then selected.

**Figure 5.**
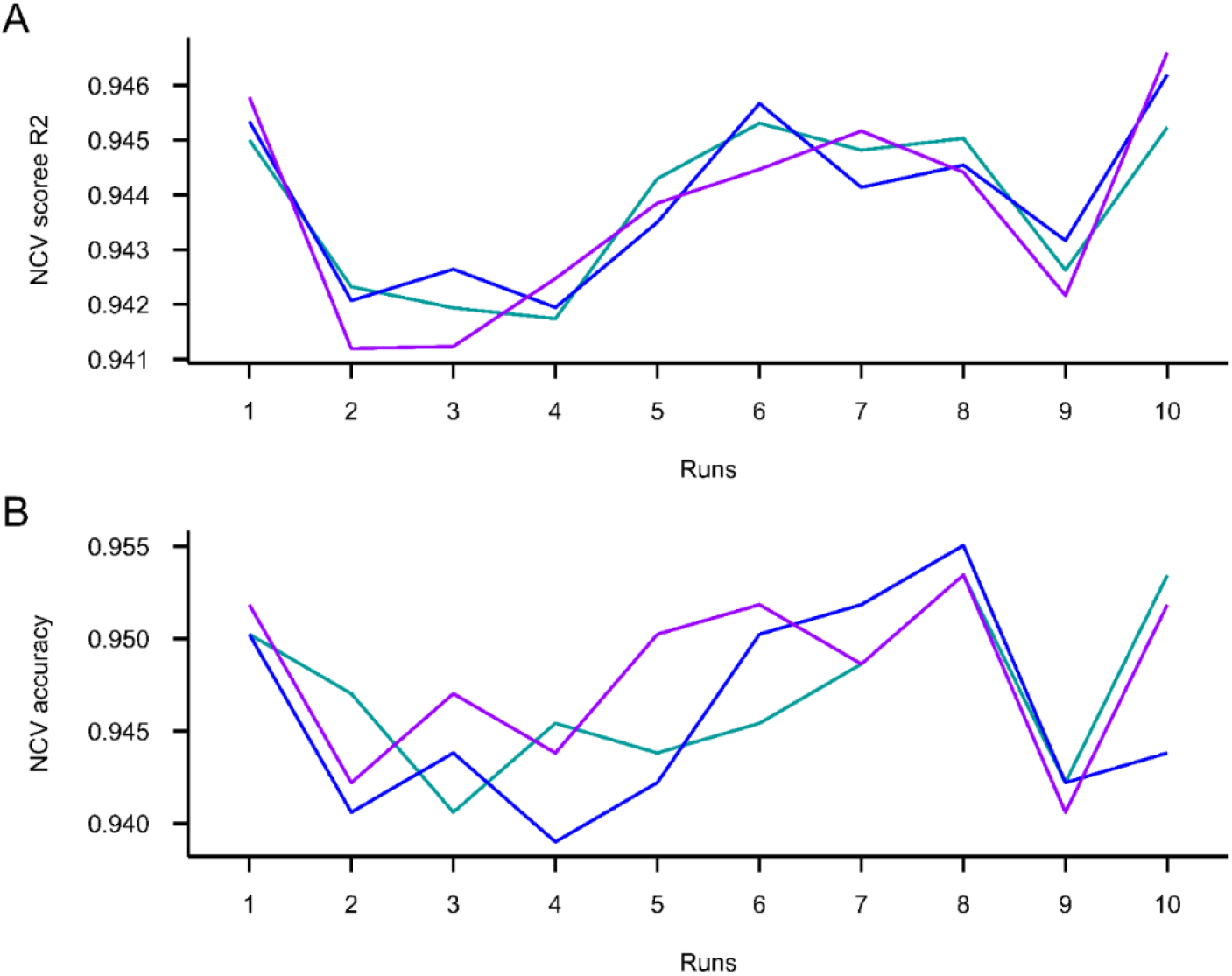
Performance comparisons of the two GB models and their aggregation across 10 runs. A. Comparison of average NCV R2 score of different models of each run. B. Comparison of average NCF accuracy of different models of each run. Gradient Boosting 1 (GB1) (green), Gradient Boosting 2 (GB2) (blue) and aggregation (purple).

Figure 6 shows the detailed results of the less performant run of all (run 9) obtained with the best model. The confusion matrix shows predictions about 623 TNBC breast cancer patients (Figure 6A). We were able to distinguish the TNBC subtypes with a high score of accuracy of 94.22%. The three TNBC subtypes had both a high sensitivity and a high specificity (Figure 6B). More particularly, patients belonging to C1 subtype were identified with a sensitivity of 97.39% and specificity of 99.57% (Figure 6B). It results from the biological characteristics of C1 which are hugely different from C2 and C3. The most incorrectly cluster assignments were in the overlap between C2 and C3 (Figure 6A and C).

**Figure 6.**
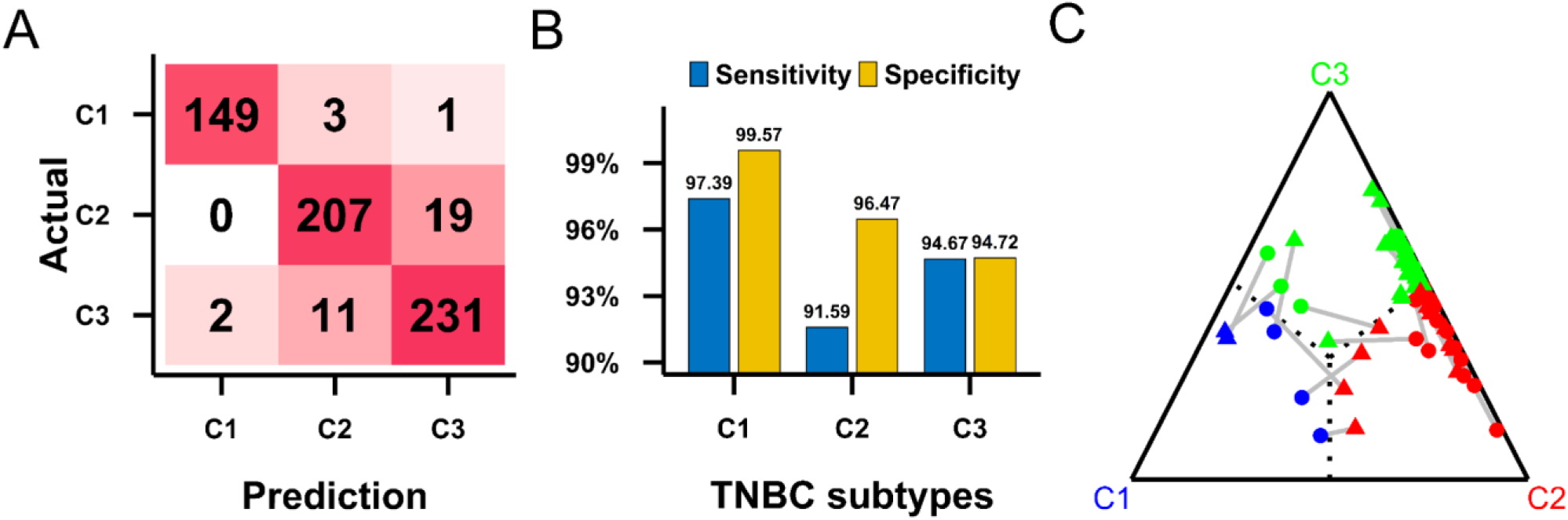
Aggregative model performance based on training data for the run 9 of NCV. A. Confusion matrix for C1, C2 and C3 TNBC subtypes. The number of correct and incorrect prediction are summarized with count and subdivided by their actual and predicted TNBC subtypes. B. Bar chart comparison of sensitivity and specificity among the three TNBC subtypes. C. Distribution of misclassified patients based on probability of belonging to TNBC subtypes. Actual probabilities (dots) were connected with prediction values (triangles) using grey segments.

This result comes as no surprise since C2 and C3 are both basal-like subtypes mainly distinguished by the direction of the immune response (that is, pro-tumorigenic in C2 and anti-tumorigenic in C3). Furthermore, we demonstrated that immunological response gradients existed between these two clusters [4]. On the contrary, biological characteristics of C1, which is a molecular apocrine cluster, are very different from those of C2 and C3.

We used this best model to predict TNBC subtypes of 70 patients of the test set. Finally, our model predicted 64 correct classifications among 70. The matrix confusion was used to summarizing the performance of the model (Figure 7A).

**Figure 7.**
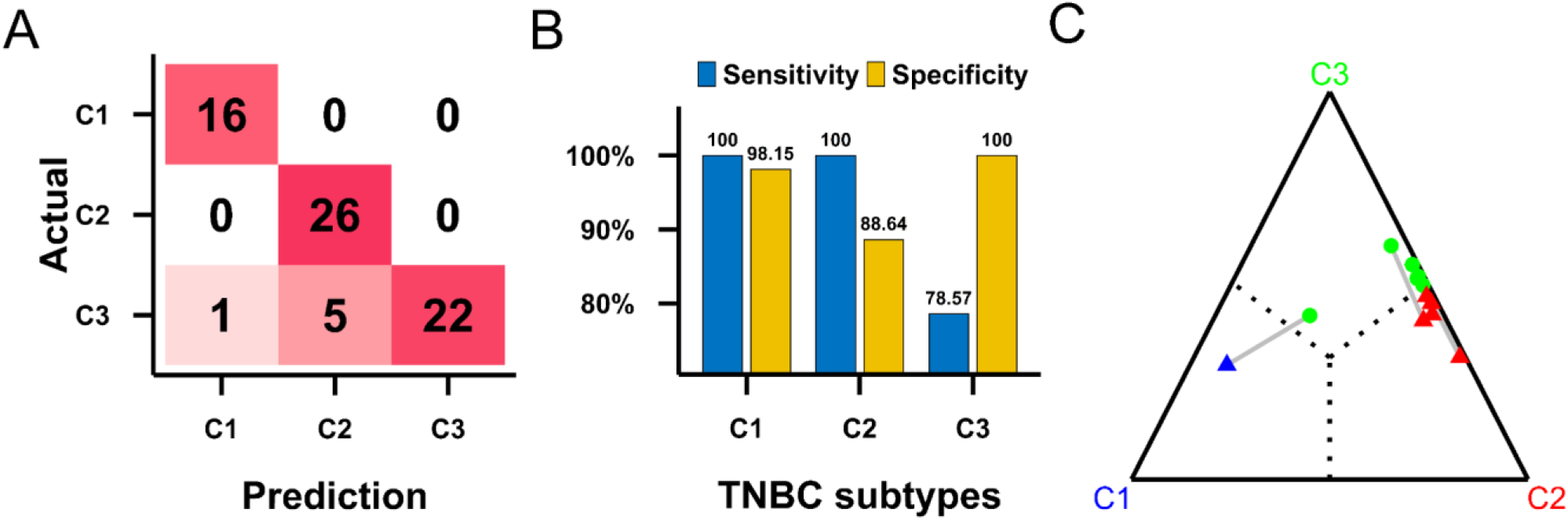
Aggregative model performance based on testing data. A. Confusion matrix for C1, C2 and C3 TNBC subtypes. The number of correct and incorrect prediction are summarized with count and subdivided by their actual and predicted TNBC subtypes. B. Bar chart comparison of sensitivity and specificity among the three TNBC subtypes. C. Distribution of misclassified patients based on probability of belonging to TNBC subtypes. Actual probabilities (dots) were connected with prediction values (triangles) using grey segments.

Overall, we can conclude that our model with aggregation of the two GB based on the List 2 of indicators is robust with a good generalization performance to predict TNBC subtypes.

## Discussion

In this work, we propose a method able to distinguish TNBC subtypes independent of external data, and with a restricted number of probes. Microarray expression studies are often compromised by the presence of batch effect and normalization step and methods. The choice to work from binary comparisons allowed us to propose a robust model which is independent of these sources of variability.

We adapt machine learning algorithms to solve a multi-target regression problem. The number of probes and the number of indicators based on these probes are reduced for both industrial and statistical performance reasons. The number of indicators was reduced from more than one billion to only 50 strongly relevant indicators, through the application of a hybrid method of variable selection on the training data (90% of full data). These indicators were generated by 36 probes. From a biological standpoint, selected probes/genes and indicators are very pertinent. These results are in total concordance with biological characteristics of the three TNBC clusters.

We found that the model with aggregation of the two GB has achieved the best NCV score performances across 10 runs of all tested models. The model’s generalization performance was approved; it generalized well to test data set. We can safely assume that it is no overfitted to the training data. Misclassified cases relate to tumors which locate at the border of fuzzy clusters, and more specially between C2 and C3. Indeed, these two basal-like clusters are very similar. The main biological trait that differentiates them is the direction of the immune response, with a decreasing pro-tumorigenic immune response from C2 to C3 and a decreasing anti-tumorigenic immune response from C3 to C2. These biological gradients create a kind of overlap between C2 and C3, which complicates prediction analysis when fuzzy clustering probabilities are close to 0.5.

In future works, we intend to adapt some approaches of structured regularizers for different types of high-dimensional problems to provide a penalization method for the problem studied in this paper [31].

## Acknowledgements

This paper was prepared in the context of the SIRIC ILIAD program supported by the French National Cancer Institute national (INCa), the Ministry of Health and the Institute for Health and Medical Research (Inserm) (SIRIC ILIAD, INCa-DGOS Inserm-12558).

## Authors’ contributions

PJ, FBA, BM and CGC conceived and designed the project. WG and HL participated in TNBC data collection. FBA, AFF, FG, TL, HL and CGC participated in the programing and statistical analyses. PJ, MC and PPJ interpreted the biological meaning of the statistical results. FBA, PJ, BM and HL wrote the manuscript. WG, PPJ and MC edited and revised the manuscript. All authors read and approved the final manuscript.

