## Supplementary data for "Development of an absolute assignment predictor for triple-negative breast cancer subtyping using machine learning approaches"

### Table S1 TNBC cohorts

| Study code | DNA chip | TNBC patients | Reference |
|---|---|---|---|
| E-MTAB-365 | Affymetrix U133P2 | 51 | (S1) |
| GSE12276 | Affymetrix U133P2 | 56 | (S2) |
| GSE18864 | Affymetrix U133P2 | 35 | (S3) |
| GSE19615 | Affymetrix U133P2 | 28 | (S4) |
| GSE21653 | Affymetrix U133P2 | 87 | (S5) |
| GSE58812 | Affymetrix U133P2 | 107 | (S6) |
| GSE76124 | Affymetrix U133P2 | 198 | (S7) |
| GSE83937 | Affymetrix U133P2 | 131 | (S8) |
| | Total | 693 | |

**Table S2 List of probes and genes selected by forward selection in function of the three probe threshold filter lists**

List 1: 0.2%; List 2: 0.4%; List 3: 0.6%.

| | **List 1** | | **List 2** | | **List 3** | |
|---|---|---|---|---|---|---|
| | **Probe set ID** | **Gene symbol** | **Probe set ID** | **Gene symbol** | **Probe set ID** | **Gene symbol** |
| 1 | 204667_at | *FOXA1* | 204667_at | *FOXA1* | 204061_at | *PRKX* |
| 2 | 204822_at | *TTK* | 204822_at | *TTK* | 204667_at | *FOXA1* |
| 3 | 205267_at | *POU2AF1* | 205267_at | *POU2AF1* | 205267_at | *POU2AF1* |
| 4 | 219497_s_at | *BCL11A* | 219497_s_at | *BCL11A* | 228214_at | *SOX6* |
| 5 | 226120_at | *TTC8* | 230538_at | *SHC4* | 205831_at | *CD2* |
| 6 | 213915_at | *NKG7* | 213915_at | *NKG7* | 219498_s_at | *BCL11A* |
| 7 | 219615_s_at | *KCNK5* | 216092_s_at | *SLC7A8* | 212985_at | *APBB2* |
| 8 | 213492_at | *COL2A1* | 1555638_a_at | *SAMSN1* | 219243_at | *GIMAP4* |
| 9 | 225968_at | *PRICKLE2* | 201951_at | *ALCAM* | 207828_s_at | *CENPF* |
| 10 | 212592_at | *JCHAIN* | 34210_at | *CD52* | 230538_at | *SHC4* |
| 11 | 209606_at | *CYTIP* | 212985_at | *APBB2* | 217480_x_at | No name |
| 12 | 219735_s_at | *TFCP2L1* | 205831_at | *CD2* | 226197_at | *AR* |
| 13 | 224428_s_at | *CDCA7* | 217148_x_at | *IGLV2-14* | 221203_s_at | *YEATS2* |
| 14 | 213260_at | *FOXC1* | 209612_s_at | *ADH1B* | 206513_at | *AIM2* |
| 15 | 226661_at | *CDCA2* | 227642_at | *TFCP2L1* | 204912_at | *IL10RA* |
| 16 | 211643_x_at | *IGKC* | 206560_s_at | *MIA* | 1554572_a_at | *SUV39H2* |
| 17 | 228969_at | *AGR2* | 221874_at | *KIAA1324* | 217281_x_at | No name |
| 18 | 227346_at | *IKZF1* | 203343_at | *UGDH* | 226248_s_at | *KIAA1324* |
| 19 | 201952_at | *ALCAM* | 231747_at | *CYSLTR1* | 1553613_s_at | *FOXC1* |
| 20 | 222891_s_at | *BCL11A* | 216401_x_at | No name | 225540_at | *MAP2* |
| 21 | 216401_x_at | No name | 205692_s_at | *CD38* | 217157_x_at | No name |
| 22 | 201124_at | *ITGB5* | 226034_at | *DUSP4* | 212830_at | MEGF9 |
| 23 | 220425_x_at | No name | 224428_s_at | *CDCA7* | 200099_s_at | No name |
| 24 | 205488_at | *GZMA* | 202274_at | *ACTG2* | 225327_at | *FAM214A* |
| 25 | 232234_at | *SLA2* | 219812_at | *PVRIG* | 219159_s_at | *SLAMF7* |
| 26 | 201468_s_at | *NQO1* | 226197_at | *AR* | 219735_s_at | *TFCP2L1* |
| 27 | 235020_at | *TAF4B* | 226661_at | *CDCA2* | 213523_at | *CCNE1* |
| 28 | 211110_s_at | *AR* | 221185_s_at | *IQCG* | 209842_at | *SOX10* |
| 29 | 236280_at | *P2RY10* | 204951_at | *RHOH* | 215946_x_at | *IGLL3P* |
| 30 | 214916_x_at | No name | 218963_s_at | *KRT23* | 219209_at | *IFIH1* |
| 31 | 213539_at | *CD3D* | 212592_at | *JCHAIN* | AFFX-HSAC07/X00351_5_at | *ACTB* |
| 32 | 227052_at | *SMIM14* | 1553613_s_at | *FOXC1* | 214053_at | *ERBB4* |
| 33 | | | 210347_s_at | *BCL11A* | 224146_s_at | *ABCC11* |
| 34 | | | 221591_s_at | *PIMREG* | 221591_s_at | *PIMREG* |
| 35 | | | 213539_at | *CD3D* | | |
| 36 | | | 204205_at | *APOBEC3G* | | |

**Table S3 List of indicators selected by forward selection in function of the three probe threshold filter lists**
List 1: 0.2%; List 2: 0.4%; List 3: 0.6%.

| | List 1 | | List 2 | | List 3 | |
|---|---|---|---|---|---|---|
| | Probe set ID | Gene symbol | Probe set ID | Gene symbol | Probe set ID | Gene symbol |
| 1 | 204667_at > 204822_at | FOXA1 > TTK | 204667_at > 204822_at | FOXA1 > TTK | 204667_at > 219498_s_at | FOXA1 > BCL11A |
| 2 | 219615_s_at > 213539_at | KCNK5 > CD3D | 212985_at > 205831_at | APBB2 > CD2 | 205831_at > 212830_at | CD2 > MEGF9 |
| 3 | 201124_at > 236280_at | ITGB5 > P2RY10 | 205267_at > 219497_s_at | POU2AF1 > BCL11A | 230538_at > 217157_x_at | SHC4 > No name |
| 4 | 205267_at > 201952_at | POU2AF1 > ALCAM | 230538_at > 34210_at | SHC4 > CD52 | 226248_s_at > 219209_at | KIAA1324 > IFIH1 |
| 5 | 219497_s_at > 211110_s_at | BCL11A > AR | 205267_at > 226197_at | POU2AF1 > AR | 228214_at > 219243_at | SOX6 > GIMAP4 |
| 6 | 212592_at > 220425_x_at | JCHAIN > No name | 204667_at > 224428_s_at | FOXA1 > CDCA7 | 207828_s_at > 226197_at | CENPF > AR |
| 7 | 219615_s_at > 228969_at | KCNK5 > AGR2 | 205831_at > 1553613_s_at | CD2 > FOXC1 | 205267_at > 212985_at | POU2AF1 > APBB2 |
| 8 | 219497_s_at > 214916_x_at | BCL11A > No name | 216401_x_at > 226661_at | No name > CDCA2 | 209842_at > 215946_x_at | SOX10 > IGLL3P |
| 9 | 209606_at > 235020_at | CYTIP > TAF4B | 216092_s_at > 204951_at | SLC7A8 > RHOH | 228214_at > 219159_s_at | SOX6 > SLAMF7 |
| 10 | 225968_at > 236280_at | PRICKLE2 > P2RY10 | 204667_at > 227642_at | FOXA1 > TFCP2L1 | 204667_at > 217281_x_at | FOXA1 > No name |
| 11 | 225968_at > 205488_at | PRICKLE2 > GZMA | 206560_s_at > 204951_at | MIA > RHOH | 219498_s_at > 219243_at | BCL11A > GIMAP4 |
| 12 | 204667_at > 222891_s_at | FOXA1 > BCL11A | 212985_at > 216401_x_at | APBB2 > No name | 205831_at > 219735_s_at | CD2 > TFCP2L1 |
| 13 | 204667_at > 224428_s_at | FOXA1 > CDCA7 | 204667_at > 219497_s_at | FOXA1 > BCL11A | 226197_at > 217157_x_at | AR > No name |
| 14 | 205267_at > 213260_at | POU2AF1 > FOXC1 | 217148_x_at > 218963_s_at | IGLV2-14 > KRT23 | 205831_at > 212985_at | CD2 > APBB2 |
| 15 | 213260_at > 201124_at | FOXC1 > ITGB5 | 226197_at > 221185_s_at | AR > IQCG | 204061_at > 212830_at | PRKX > MEGF9 |
| 16 | 213492_at > 236280_at | COL2A1 > P2RY10 | 204822_at > 212592_at | TTK > JCHAIN | 205267_at > 221203_s_at | POU2AF1 > YEATS2 |
| 17 | 226120_at > 216401_x_at | TTC8 > No name | 203343_at > 221185_s_at | UGDH > IQCG | 212985_at > 217480_x_at | APBB2 > No name |
| 18 | 219615_s_at > 201124_at | KCNK5 > ITGB5 | 201951_at > 205692_s_at | ALCAM > CD38 | 219209_at > AFFX-HSAC07/X00351_5_at | IFIH1 > ACTB |
| 19 | 205267_at > 219735_s_at | POU2AF1 > TFCP2L1 | 205692_s_at > 221591_s_at | CD38 > PIMREG | 212985_at > 219159_s_at | APBB2 > SLAMF7 |
| 20 | 212592_at > 222891_s_at | JCHAIN > BCL11A | 206560_s_at > 212592_at | MIA > JCHAIN | 217157_x_at > 219735_s_at | No name > TFCP2L1 |
| 21 | 213915_at > 219615_s_at | NKG7 > KCNK5 | 201951_at > 34210_at | ALCAM > CD52 | 225327_at > 219209_at | FAM214A > IFIH1 |
| 22 | 220425_x_at > 213539_at | No name > CD3D | 231747_at > 226034_at | CYSLTR1 > DUSP4 | 204667_at > 221591_s_at | FOXA1 > PIMREG |
| 23 | 226661_at > 211110_s_at | CDCA2 > AR | 209612_s_at > 204205_at | ADH1B > APOBEC3G | 219243_at > 230538_at | GIMAP4 > SHC4 |
| 24 | 226120_at > 213539_at | TTC8 > CD3D | 205831_at > 203343_at | CD2 > UGDH | 204912_at > 225540_at | IL10RA > MAP2 |
| 25 | 204667_at > 211643_x_at | FOXA1 > IGKC | 227642_at > 204205_at | TFCP2L1 > APOBEC3G | 204061_at > 204667_at | PRKX > FOXA1 |
| 26 | 201952_at > 216401_x_at | ALCAM > No name | 213915_at > 221874_at | NKG7 > KIAA1324 | 226197_at > 206513_at | AR > AIM2 |
| 27 | 228969_at > 211110_s_at | AGR2 > AR | 224428_s_at > 226197_at | CDCA7 > AR | 204667_at > 207828_s_at | FOXA1 > CENPF |
| 28 | 219497_s_at > 211643_x_at | BCL11A > IGKC | 216401_x_at > 202274_at | No name > ACTG2 | 221203_s_at > 204912_at | YEATS2 > IL10RA |
| 29 | 213492_at > 216401_x_at | COL2A1 > No name | 206560_s_at > 213539_at | MIA > CD3D | 221203_s_at > 217157_x_at | YEATS2 > No name |
| 30 | 219615_s_at > 205488_at | KCNK5 > GZMA | 204822_at > 212985_at | TTK > APBB2 | 206513_at > 226248_s_at | AIM2 > KIAA1324 |
| 31 | 205267_at > 219497_s_at | POU2AF1 > BCL11A | 204667_at > 226661_at | FOXA1 > CDCA2 | 219243_at > 226197_at | GIMAP4 > AR |
| 32 | 219497_s_a t> 201124_at | BCL11A > ITGB5 | 230538_at > 209612_s_at | SHC4 > ADH1B | 219498_s_at > 206513_at | BCL11A > AIM2 |
| 33 | 201952_at > 232234_at | ALCAM > SLA2 | 219497_s_at > 227642_at | BCL11A > TFCP2L1 | 226197_at > 209842_at | AR > SOX10 |
| 34 | 201124_at > 205488_at | ITGB5 > GZMA | 231747_at > 210347_s_at | CYSLTR1 > BCL11A | 219243_at > 209842_at | GIMAP4 > SOX10 |
| 35 | 204667_at > 235020_at | FOXA1 > TAF4B | 226197_at > 218963_s_at | AR > KRT23 | 205831_at > 219498_s_at | CD2 > BCL11A |
| 36 | 226120_at > 214916_x_at | TTC8 > No name | 213915_at > 204205_at | NKG7 > APOBEC3G | 205267_at > AFFX-HSAC07/X00351_5_at | POU2AF1 > ACTB |
| 37 | 219735_s_at > 227346_at | TFCP2L1 > IKZF1 | 226197_at > 1553613_s_at | AR > FOXC1 | 205831_at > 225327_at | CD2 > FAM214A |
| 38 | 225968_at > 226661_at | PRICKLE2 > CDCA2 | 209612_s_at > 216401_x_at | ADH1B > No name | 226197_at > 1553613_s_at | AR > FOXC1 |
| 39 | 226120_at > 236280_at | TTC8 > P2RY10 | 205267_at > 221874_at | POU2AF1 > KIAA1324 | 205831_at > 221591_s_at | CD2 > PIMREG |
| 40 | 219497_s_at > 205488_at | BCL11A > GZMA | 204822_at > 205831_at | TTK > CD2 | 230538_at > 217281_x_at | SHC4 > No name |
| 41 | 205267_at > 235020_at | POU2AF1 > TAF4B | 204822_at > 231747_at | TTK > CYSLTR1 | 230538_at > 225540_at | SHC4 > MAP2 |
| 42 | 213915_at > 201124_at | NKG7 > ITGB5 | 205831_at > 209612_s_at | CD2 > ADH1B | 209842_at > 214053_at | SOX10 > ERBB4 |
| 43 | 219497_s_at > 225968_at | BCL11A > PRICKLE2 | 226034_at > 226661_at | DUSP4 > CDCA2 | 221203_s_at > 225327_at | YEATS2 > FAM214A |
| 44 | 235020_at > 213539_at | TAF4B > CD3D | 203343_at > 216401_x_at | UGDH > No name | 221203_s_at > 219159_s_at | YEATS2 > SLAMF7 |
| 45 | 209606_at > 201468_s_at | CYTIP > NQO1 | 206560_s_at > 204205_at | MIA > APOBEC3G | 212830_at > 219209_at | MEGF9 > IFIH1 |
| 46 | 226120_at > 209606_at | TTC8 > CYTIP | 205831_at > 221591_s_at | CD2 > PIMREG | 217480_x_at > 225327_at | No name > FAM214A |
| 47 | 213915_at > 213492_at | NKG7 > COL2A1 | 205267_at > 203343_at | POU2AF1 > UGDH | 212985_at > 217281_x_at | APBB2 > No name |
| 48 | 201468_s_at > 213539_at | NQO1 > CD3D | 203343_at > 218963_s_at | UGDH > KRT23 | 212830_at > 225327_at | MEGF9 > FAM214A |
| 49 | 211110_s_at > 236280_at | AR > P2RY10 | 230538_at > 226197_at | SHC4 > AR | 230538_a t > 219159_s_at | SHC4 > SLAMF7 |
| 50 | 204822_at > 211110_s_at | TTK > AR | 231747_at > 226197_at | CYSLTR1 > AR | 217281_x_at > 225540_at | No name > MAP2 |

**Table S4 Comparisons of different models using different list of indicators and 10 runs of NCV in terms of R2 score**
Random Forest (RF), Gradient Boosting 1 (GB1), Gradient Boosting 2 (GB2), XGBoost 1 (XGB1) and XGBoost 2 (XGB2).

| | List 1 | | | | | List 2 | | | | | List3 | | | | |
|---|---|---|---|---|---|---|---|---|---|---|---|---|---|---|---|
| **Models** | **RF** | **GB1** | **GB2** | **XGB1** | **XGB2** | **RF** | **GB1** | **GB2** | **XGB1** | **XGB2** | **RF** | **GB1** | **GB2** | **XGB1** | **XGB2** |
| Run 1 | 0.9264 | 0.9303 | 0.9306 | 0.9201 | 0.9205 | 0.9328 | 0.9450 | 0.9453 | 0.9397 | 0.9353 | 0.9362 | 0.9430 | 0.9390 | 0.9366 | 0.9339 |
| Run 2 | 0.9258 | 0.9335 | 0.9342 | 0.9195 | 0.9190 | 0.9271 | 0.9421 | 0.9412 | 0.9377 | 0.9329 | 0.9370 | 0.9431 | 0.9418 | 0.9372 | 0.9369 |
| Run 3 | 0.9267 | 0.9326 | 0.9336 | 0.9230 | 0.9214 | 0.9297 | 0.9412 | 0.9419 | 0.9346 | 0.9303 | 0.9325 | 0.9422 | 0.9426 | 0.9306 | 0.9265 |
| Run 4 | 0.9278 | 0.9354 | 0.9361 | 0.9275 | 0.9234 | 0.9320 | 0.9417 | 0.9419 | 0.9376 | 0.9355 | 0.9332 | 0.9426 | 0.9418 | 0.9332 | 0.9289 |
| Run 5 | 0.9289 | 0.9344 | 0.9345 | 0.9292 | 0.9272 | 0.9330 | 0.9435 | 0.9439 | 0.9411 | 0.9370 | 0.9341 | 0.9436 | 0.9415 | 0.9347 | 0.9326 |
| Run 6 | 0.9261 | 0.9322 | 0.9325 | 0.9234 | 0.9241 | 0.9328 | 0.9445 | 0.9453 | 0.9357 | 0.9341 | 0.9382 | 0.9448 | 0.9451 | 0.9407 | 0.9374 |
| Run 7 | 0.9267 | 0.9351 | 0.9358 | 0.9235 | 0.9212 | 0.9335 | 0.9448 | 0.9441 | 0.9400 | 0.9328 | 0.9361 | 0.9437 | 0.9431 | 0.9384 | 0.9367 |
| Run 8 | 0.9266 | 0.9316 | 0.9330 | 0.9244 | 0.9212 | 0.9344 | 0.9445 | 0.9444 | 0.9408 | 0.9385 | 0.9349 | 0.9431 | 0.9433 | 0.9363 | 0.9357 |
| Run 9 | 0.9276 | 0.9328 | 0.9329 | 0.9243 | 0.9233 | 0.9334 | 0.9422 | 0.9426 | 0.9371 | 0.9335 | 0.9331 | 0.9408 | 0.9411 | 0.9365 | 0.9334 |
| Run 10 | 0.9281 | 0.9325 | 0.9336 | 0.9258 | 0.9255 | 0.9351 | 0.9452 | 0.9462 | 0.9403 | 0.9403 | 0.9393 | 0.9439 | 0.9426 | 0.9372 | 0.9334 |
| Mean | 0.9271 | 0.9330 | 0.9337 | 0.9241 | 0.9227 | 0.9324 | 0.9435 | 0.9437 | 0.9385 | 0.9350 | 0.9355 | 0.9431 | 0.9422 | 0.9361 | 0.9335 |
| Median | 0.9267 | 0.9327 | 0.9336 | 0.9239 | 0.9223 | 0.9329 | 0.9440 | 0.9440 | 0.9387 | 0.9347 | 0.9355 | 0.9431 | 0.9422 | 0.9366 | 0.9337 |

**Table S5 Comparisons of different models using different list of indicators and 10 runs of NCV in terms of accuracy**
Random Forest (RF), Gradient Boosting 1 (GB1), Gradient Boosting 2 (GB2), XGBoost 1 (XGB1) and XGBoost 2 (XGB2).

| | List 1 | | | | | List 2 | | | | | List3 | | | | |
|---|---|---|---|---|---|---|---|---|---|---|---|---|---|---|---|
| **Models** | **RF** | **GB1** | **GB2** | **XGB1** | **XGB2** | **RF** | **GB1** | **GB2** | **XGB1** | **XGB2** | **RF** | **GB1** | **GB2** | **XGB1** | **XGB2** |
| Run 1 | 0.9278 | 0.9262 | 0.9278 | 0.9230 | 0.9101 | 0.9470 | 0.9502 | 0.9502 | 0.9454 | 0.9406 | 0.9374 | 0.9390 | 0.9326 | 0.9342 | 0.9278 |
| Run 2 | 0.9326 | 0.9358 | 0.9406 | 0.9294 | 0.9278 | 0.9358 | 0.9406 | 0.9422 | 0.9374 | 0.9278 | 0.9326 | 0.9454 | 0.9390 | 0.9406 | 0.9406 |
| Run 3 | 0.9294 | 0.9294 | 0.9326 | 0.9262 | 0.9213 | 0.9390 | 0.9470 | 0.9406 | 0.9358 | 0.9294 | 0.9422 | 0.9470 | 0.9454 | 0.9294 | 0.9294 |
| Run 4 | 0.9326 | 0.9310 | 0.9294 | 0.9262 | 0.9197 | 0.9422 | 0.9454 | 0.9390 | 0.9342 | 0.9342 | 0.9454 | 0.9454 | 0.9438 | 0.9406 | 0.9294 |
| Run 5 | 0.9342 | 0.9326 | 0.9310 | 0.9326 | 0.9326 | 0.9454 | 0.9422 | 0.9502 | 0.9470 | 0.9486 | 0.9454 | 0.9422 | 0.9374 | 0.9310 | 0.9326 |
| Run 6 | 0.9310 | 0.9310 | 0.9294 | 0.9230 | 0.9230 | 0.9470 | 0.9518 | 0.9454 | 0.9422 | 0.9374 | 0.9406 | 0.9454 | 0.9454 | 0.9438 | 0.9310 |
| Run 7 | 0.9262 | 0.9342 | 0.9326 | 0.9310 | 0.9262 | 0.9502 | 0.9486 | 0.9518 | 0.9454 | 0.9486 | 0.9438 | 0.9454 | 0.9454 | 0.9406 | 0.9278 |
| Run 8 | 0.9310 | 0.9262 | 0.9326 | 0.9294 | 0.9262 | 0.9438 | 0.9551 | 0.9535 | 0.9470 | 0.9454 | 0.9470 | 0.9486 | 0.9486 | 0.9438 | 0.9390 |
| Run 9 | 0.9406 | 0.9310 | 0.9294 | 0.9406 | 0.9294 | 0.9518 | 0.9406 | 0.9422 | 0.9438 | 0.9422 | 0.9342 | 0.9390 | 0.9374 | 0.9310 | 0.9342 |
| Run 10 | 0.9278 | 0.9310 | 0.9246 | 0.9262 | 0.9197 | 0.9518 | 0.9535 | 0.9438 | 0.9470 | 0.9502 | 0.9454 | 0.9454 | 0.9470 | 0.9390 | 0.9358 |
| Mean | 0.9313 | 0.9308 | 0.9310 | 0.9287 | 0.9236 | 0.9454 | 0.9475 | 0.9459 | 0.9425 | 0.9404 | 0.9414 | 0.9443 | 0.9422 | 0.9374 | 0.9327 |
| Median | 0.9310 | 0.9310 | 0.9302 | 0.9278 | 0.9246 | 0.9462 | 0.9478 | 0.9446 | 0.9446 | 0.9414 | 0.9430 | 0.9454 | 0.9446 | 0.9398 | 0.9318 |

**Figure S1 Projection of seven TNBC cohorts in the first PCA plane**

(A) E-MTAB-365 patients (n = 51, [blue]), GSE12276 patients (n = 56, [red]), GSE18864 patients (n = 35, [green]), GSE19615 patients (n = 28, [purple]), GSE21653 patients (n = 87, [hot pink]), GSE58812 patients (n = 107, [orange]), GSE76124 patients (n = 198, [gray]) and GSE83937 patients (n = 131, [black]).

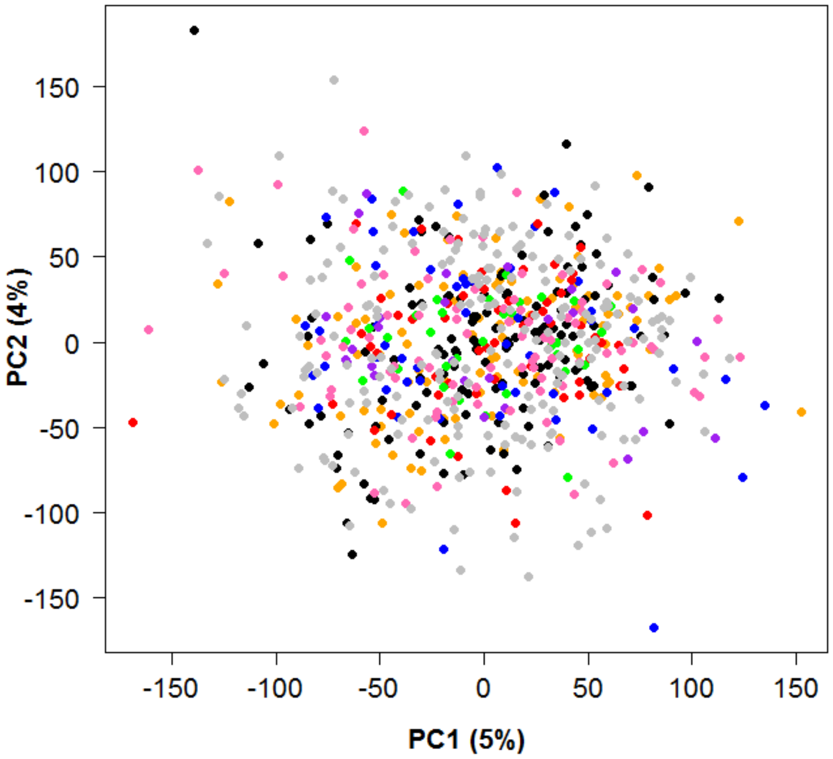

**Figure S2 Fuzzy clustering of 693 TNBC patients**

Distribution of patients based on probability of belonging to cluster: C1, n = 169 (blue); C2, n = 252 (red); C3, n = 272 (green). Each vertex of the triangle represents a cluster and each point represents a patient, placed as the barycenter of the triangle, weights being the probabilities of belonging to each of the clusters. The closer a point is to one of the vertices, the greater is the probability of the patient to belong to the corresponding cluster.

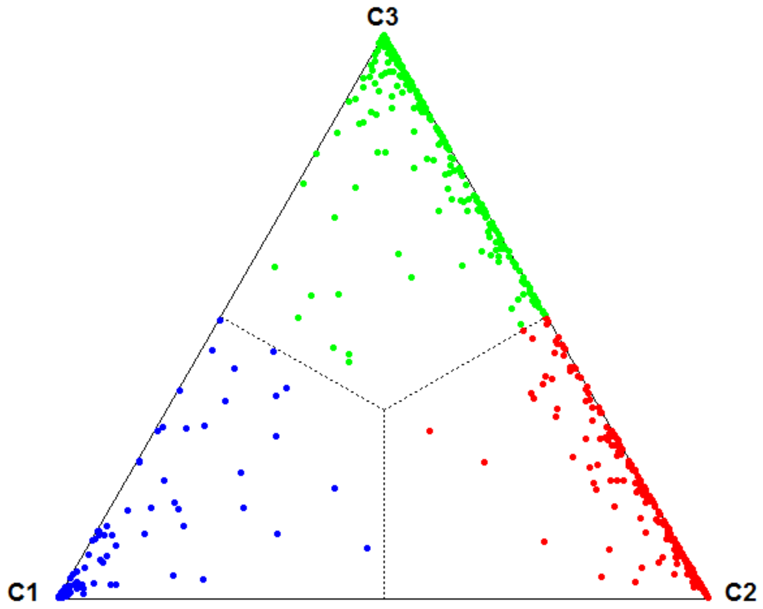

**Figure S3 Barplots of the 50 indicators of list 1, which are significantly related to the discrimination of TNBC subtypes**
They display the proportion of patients for those the value of the indicator is equal to 1 (Green), or to 0 (Red) within the TNBC subtypes.

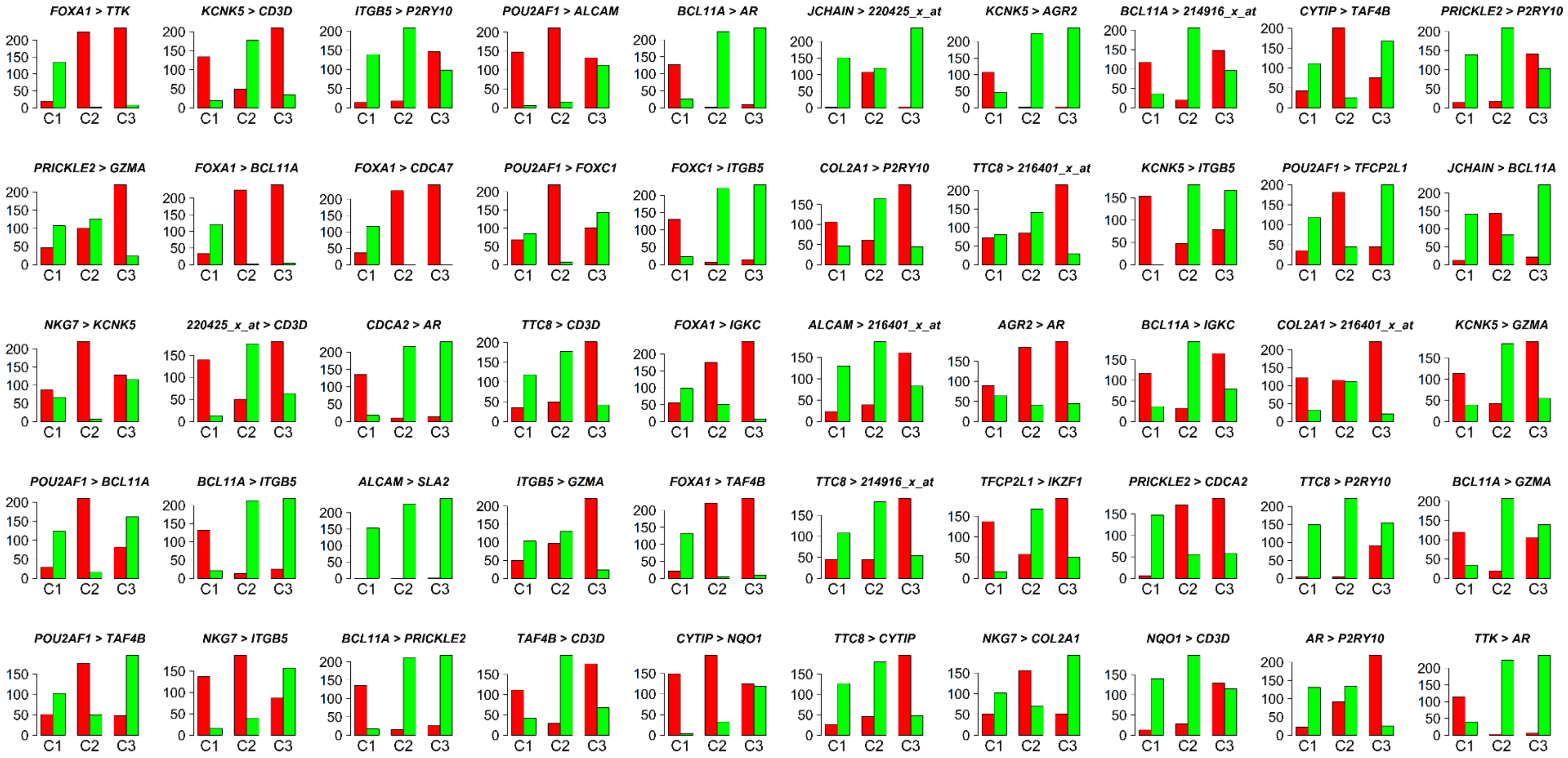

**Figure S4 Barplots of the 50 indicators of list 2, which are significantly related to the discrimination of TNBC subtypes**

They display the proportion of patients for those the value of the indicator is equal to 1 (Green), or to 0 (Red) within the TNBC subtypes.

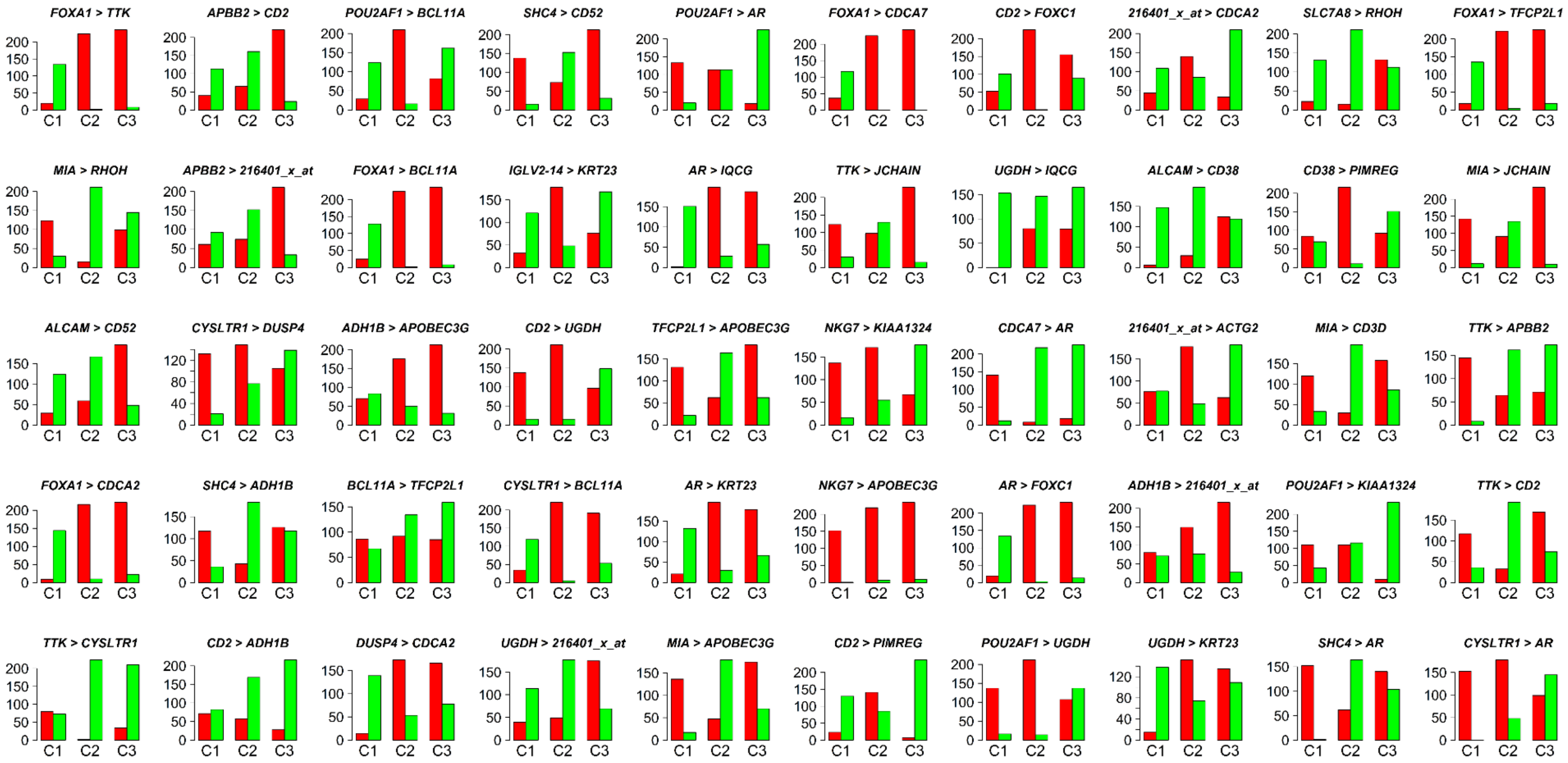

**Figure S5 Barplots of the 50 indicators of list 3, which are significantly related to the discrimination of TNBC subtypes**
They display the proportion of patients for those the value of the indicator is equal to 1 (Green), or to 0 (Red) within the TNBC subtypes.

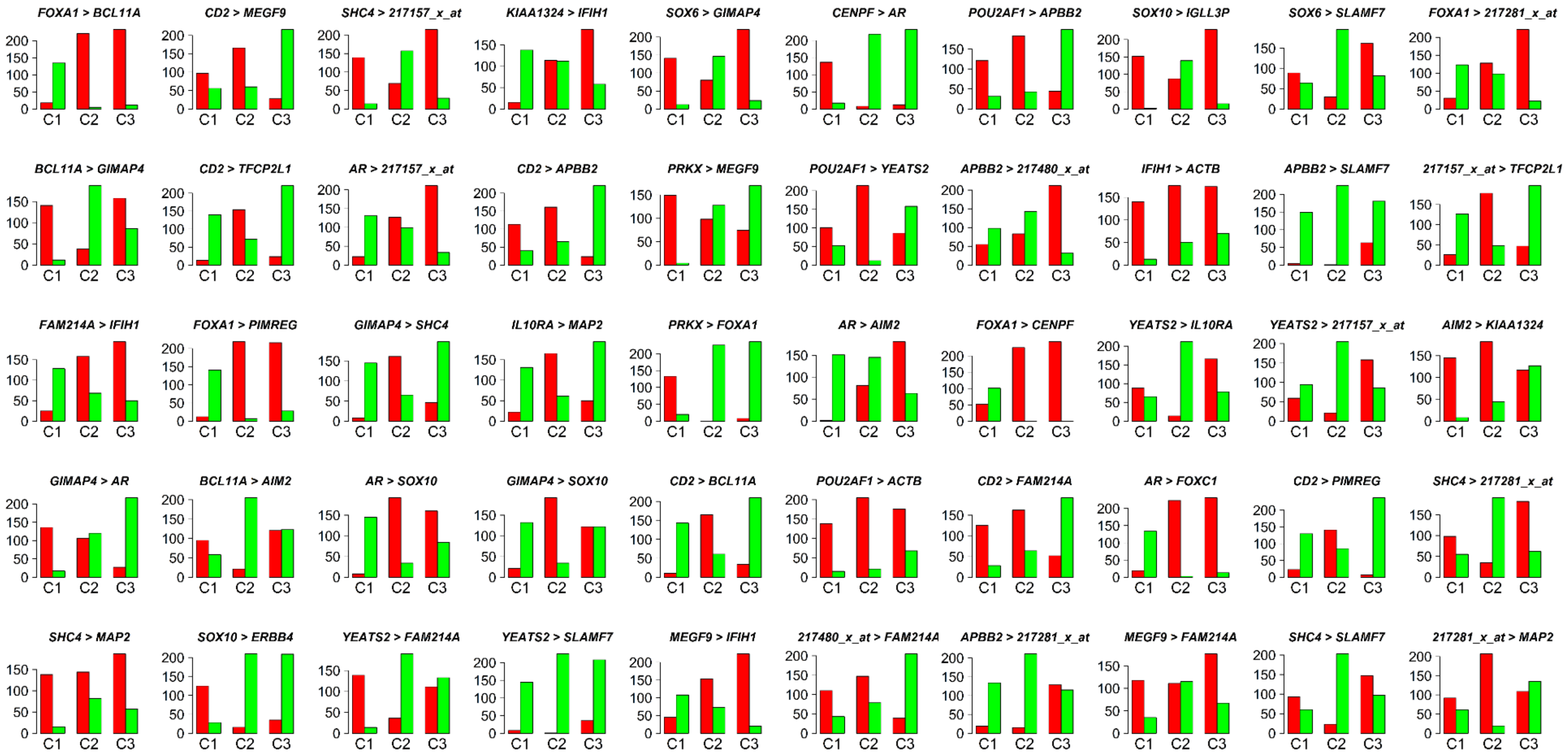
